# LS^x^: Automated reduction of gene-specific lineage evolutionary rate heterogeneity for multi-gene phylogeny inference

**DOI:** 10.1101/220053

**Authors:** Carlos J. Rivera-Rivera, Juan I. Montoya-Burgos

## Abstract

**Motivation:** LS^3^ is a recently published algorithm to reduce lineage evolutionary rate heterogeneity, a condition that can produce inference artifacts in molecular phylogenetics. The LS^3^ scripts are Linux-specific and the criterion to reduce lineage rate heterogeneity can be too stringent in datasets with both very long and very short branches.

**Results:** LS^x^ is a multi-platform user-friendly R script that performs the LS^3^ algorithm, and has added features in order to make better lineage rate calculations. In addition, we developed and implemented an alternative version of the algorithm, LS^4^, which reduces lineage rate heterogeneity not only by detecting branches that are too long but also branches that are too short, resulting in less stringent data filtering.

**Availability:** The LS^x^ script LSx_v.1.1.R and the user manual are available for download at: https://genev.unige.ch/research/laboratory/Juan-Montoya

## Introduction

We recently showed that biases emerging from evolutionary rate heterogeneity among lineages in multi-gene phylogenies can be reduced with a sequence data subselection algorithm to the point of uncovering the true phylogenetic signal (Rivera-Rivera and Montoya-Burgos 2016).

In that study, we presented an algorithm called Locus Specific Sequence Subsampling (LS^3^), which reduces lineage evolutionary rate heterogeneity gene-by-gene in multi-gene datasets. For each gene alignment, LS^3^ implements a likelihood ratio test (LRT) (Felsenstein 1981) between two models. One model assumes equal rates of evolution among all ingroup lineages (single rate model) and the other model assumes that three (or more), user-defined ingroup lineages have their own independent rate of evolution (multiple rates model). If the multiple rates model fits the data significantly better than the single rate model, the branch lengths of the phylogeny are estimated, the fastest-evolving sequence is removed, and the new reduced gene sequence dataset is tested again with the LRT. This process is iterated until a set of ingroup taxa is found whose lineage evolutionary rates can be explained equally well by the single rate model or the multiple rates model. The fast-evolving sequences that had to be removed from each gene alignment to reach this point are flagged as potentially problematic due to their higher evolutionary rate, and gene datasets which never reached this point are flagged as a whole as potentially problematic (Rivera-Rivera and Montoya-Burgos 2016). LS^3^ proved to be effective in reducing long branch attraction (LBA) artifacts in simulated nucleotide data and in two biological multi-gene datasets (one in nucleotides, and one in amino acids), and its potential to reduce phylogenetic biases has been recognized by several other authors (Cruaud and Rasplus 2016; Suh 2016; Bleidorn 2017).

Concomitantly to the publication of the LS^3^ algorithm, a set of bash scripts were made available to perform these tasks automatically (http://genev.unige.ch/en/users/Juan-Montoya/unit). While these bash scripts are effective, they only run in Linux systems, and the layout and programming is not entirely user-friendly. This prompted us to produce a new, re-programmed version of the LS^3^ algorithm which is user-friendly, contains important new features, and can be used across platforms. For this new implementation we also developed a second data subselection algorithm based on LS^3^, called “LS^3^ *supplement*” (LS^4^) which subselects sequences for lineage evolutionary rate homogeneity by not only removing very long branches, but also very short ones.

## LS^x^ Description

The new script is called LS^x^, is entirely written in R (R Core Team 2016), and uses PAML (Yang 2007) and the R packages *ape* (Paradis et al. 2004; Popescu et al. 2012) and *adephylo* (Jombart et al. 2010). If PAML, R, the R packages and Rscript are installed and functional, the script runs regardless of the platform, with all parameters given in a single raw text control file. As with the bash implementation of LS^3^, LS^x^ reads alignments in PHYLIP format, but unlike the former, it produces for each gene a version of the alignment including only the sequences that satisfied the lineage rate homogeneity criterion. In LS^x^, the best model of sequence evolution can be given for each gene (in the original implementation only a single model could be selected for all genes), thus improving the accuracy of the branch length estimations and likelihood calculations under PAML. In addition, the users can select more than three lineages of interest for the lineage evolutionary rate heterogeneity test (Fig. 1*a,b*), while the bash implementation was hardcoded to work with only three lineages of interest.

**Figure 1.**
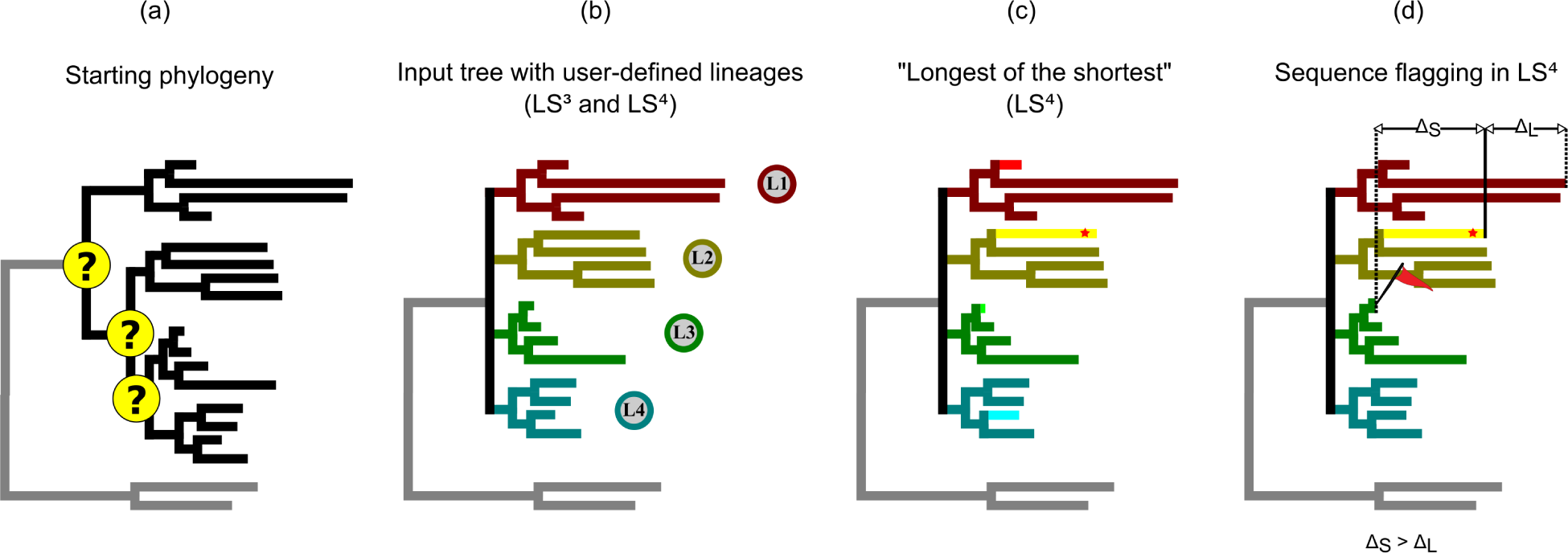
Schematic representation of the procedure for flagging sequences in LS^4^. In (*a*), the general phylogeny for this group of taxa, with the nodes in question highlighted. For for LS^3^ and LS^4^, an input tree is given in which the nodes into question are collapsed, and the lineages involved are identified (*b*). In (*c*), LS^4^ identifies the shortest branch for each clade (highlighted), and then identifies the longest among them (red star). The sequence to be removed in each iteration of LS^4^ is the one with the tip furthest from the tip of the glongest of the shortest branch (*d*), resulting in the flagging of both extremely long and extremely short branches.

## LS^3^: an LS^4^ Supplement

Within LS^x^ we also implemented LS^4^, a second data subselection algorithm that is optimized for datasets in which not only extremely long branches but also extremely short branches are present in the starting phylogenetic tree. While both LS^3^ and LS^4^ follow the same general algorithm, they differ in the criterion for choosing the sequence to be removed in the sequence subselection steps. Under LS^3^, the fastest-evolving of the ingroup sequences is removed in each iteration, as determined by its calculated sum of branch length (SBL), starting from the stem of the ingroup (Fig. 2*a*). This can lead to the flagging of too much data in a dataset containing extremely slow-evolving sequences (Fig. 2*b*). In such cases, under LS^3^, not only the fast-evolving sequences will be removed, but also the sequences with intermediate evolutionary rates which are still evolving “too fast” relative to the extremely slow-evolving ones (Fig. 2*b*). While this approach is not incorrect, simply very stringent, it can lead to the flagging of too many sequences in certain datasets and to poorly-resolved phylogenies when concatenating and analyzing the remaining data.

**Figure 2.**
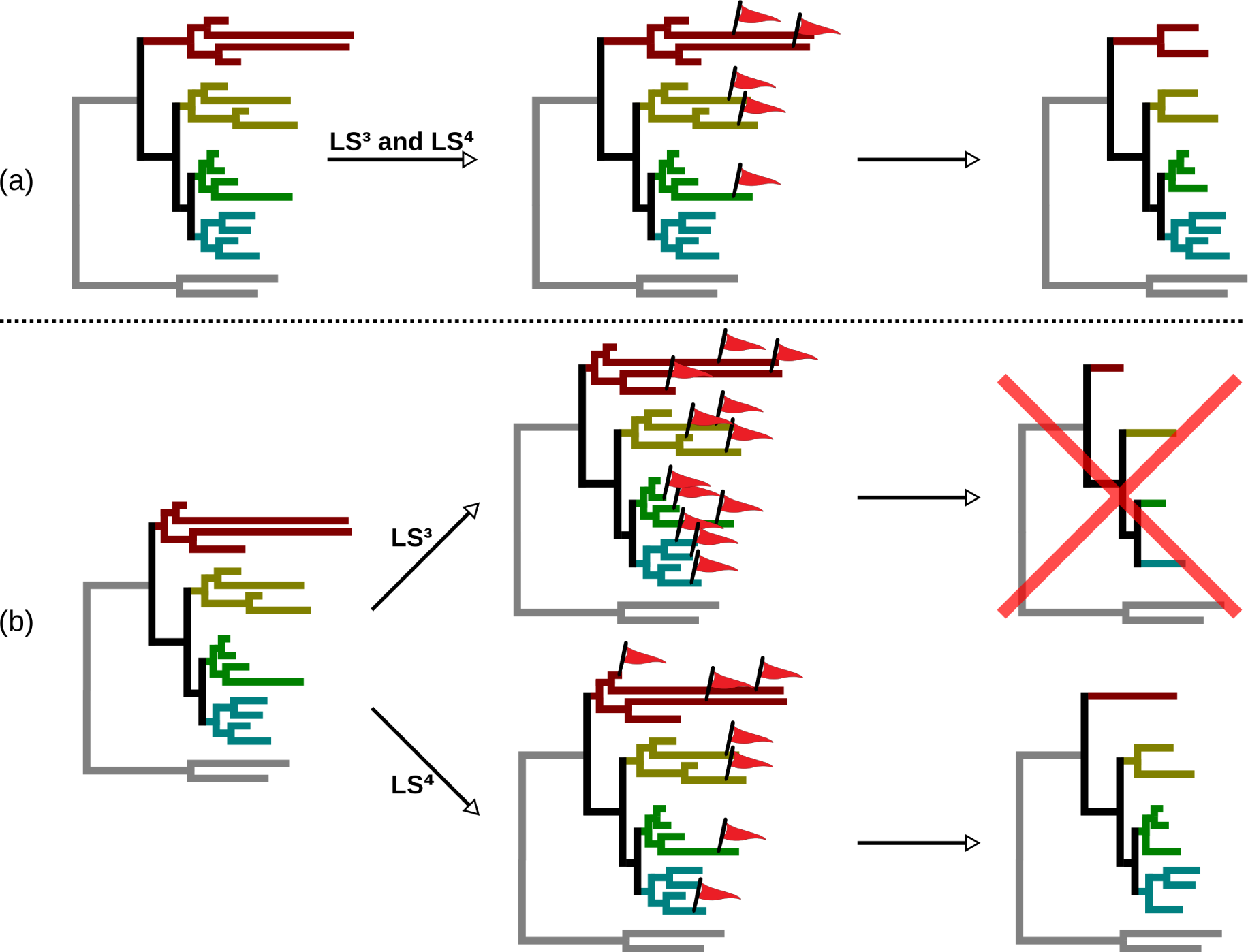
A schematic representation of the different ways that LS^3^ and LS^4^ reach lineage rate homogeneity for a given gene sequence dataset. In (*a*), a dataset with a general rate homogeneity with the exception of several faster evolving branches. In this case, both methods will flag the faster evolving sequences, and reach the same taxon subset with homogeneous rates of evolution. In (*b*), a dataset with general lineage rate heterogeneity, and with a very short branch (the top branch of that tree). In such a case, LS^3^ will remove all of the faster evolving sequences, until only the slowest sequence of each clade of interest remains. At this point, that gene sequence dataset is flagged completely as problematic because lineage rate heterogeneity is still too strong. In contrast, LS^4^ will remove the faster evolving sequences and also the slowest one, thus reaching lineage rate homogeneity for this dataset.

In order to allow for the inclusion of more data while still reaching lineage rate homogeneity, LS^4^ employs a different criterion which considers both too fast- and too slow-evolving sequences for removal. Under LS^4^, when the SBLs for all ingroup sequences of a given gene are calculated, they are grouped by the user-defined lineage of interest to which they belong. The shortest branch of each lineage of interest is identified, and then the longest among them across all ingroup lineages (“the longest of the shortest”, see Fig. 1*c*) is picked as a benchmark. Because in both LS^3^ and LS^4^ each lineage of interest has to be represented by a minimum of species (defined by the user), this “longest of the shortest” branch represents the slowest evolutionary rate at which all lineages could converge. Then, in the sequence subselection steps, the sequence to be removed is the ingroup sequence for which the absolute value of the difference between its SBL and the SBL of the benchmark sequence is the largest. In other words, for a given gene, the sequence removed in each iteration of LS^4^ is that which produces the tip furthest from the benchmark, be it faster- or slower-evolving (Fig. 1*d*). We are confident that the criterion used in LS^4^ will result in less genes flagged completely (as compared to LS^3^) when extremely slow-evolving sequences are present (Fig. 2*b*).

## Acknowledgements

We are deeply thankful to Jose Nunes for his help and suggestions during the programming of LS^x^ in R, and Joe Felsenstein for discussions about the subselection criterion used in the LS^4^ algorithm. This research was funded by the Swiss National Science Foundation (grant 31003A_141233 to JIMB) and the Institute for Genetics and Genomics in Geneva (iGE3).

